# CasanovoGUI: a cross-platform desktop application for deep learning-based *de novo* peptide sequencing with Casanovo

**DOI:** 10.64898/2026.07.11.737889

**Authors:** Bo Wen, Kai Li, Michael Riffle, Michael J. MacCoss, Wout Bittremieux, William Stafford Noble

## Abstract

*De novo* peptide sequencing detects peptides directly from tandem mass spectra without a protein sequence database, and deep learning has substantially advanced its performance. Casanovo, one such widely used model, is distributed as a Python command-line program. Consequently, installation, GPU and dependency configuration, and manual parameterization can be challenging for many bench scientists and are a recurring source of errors. Interpreting and validating the resulting predictions poses a further challenge. We present CasanovoGUI, an open-source Java-based desktop application that makes all of Casanovo’s main analysis functions available through a point-and-click interface on Windows, macOS, and Linux. On first use, CasanovoGUI automatically installs a private Python environment and Casanovo with a GPU-matched build, requiring no prior software setup. The GUI provides access to Casanovo’s analysis functions and configuration parameters, streams live progress, and integrates results interpretation: annotated spectra with per-residue confidence scores in the PDV viewer, and mismatch-tolerant mapping of *de novo* peptides back to a reference proteome. CasanovoGUI is available at https://github.com/Noble-Lab/CasanovoGUI.

## Introduction

*De novo* peptide sequencing methods infer a peptide’s amino acid sequence directly from a corresponding tandem mass spectrum, without reference to a protein sequence database. *De novo* methods are indispensable whenever the peptide of interest is unlikely to be present in the reference database used for a conventional database search—for example in antibody and immunopeptidome characterization, metaproteomics, and the discovery of variant peptides. ^1^ Recently, as in many other fields, ^2^ *de novo* sequencing has advanced rapidly with deep learning, and methods that learn to map directly from spectrum to sequence ^1,3^ have largely supplanted earlier graph- and probabilistic-network approaches. ^4,5^ Among recent methods, Casanovo ^6^ is a widely used, high-performance model that treats spectrum annotation as a sequence-to-sequence translation problem with a transformer architecture, ^7^ and that additionally supports database search using the same scoring model. ^8^

Currently, Casanovo is distributed solely as a Python command-line program, which creates a substantial practical barrier for many of the bench scientists who generate and interpret proteomics tandem mass spectrometry (MS/MS) data. Running even a single analysis requires the user to install a compatible Python interpreter, create an isolated software environment, install Casanovo together with a deep-learning stack such as PyTorch and the appropriate CUDA libraries, and then compose command lines and edit a parameter configuration file by hand. These steps are both time-consuming and error-prone. For example, incompatible native libraries can cause Casanovo to fail at startup with opaque memory-access errors, and routine package updates can silently replace a GPU-enabled installation with a much slower CPU-only build. For users without programming experience, these obstacles may prevent adoption altogether; for experienced users, they still consume time and sometimes create hard-to-diagnose failures.

A second barrier concerns the interpretation of *de novo* sequencing results. Unlike database search, *de novo* sequencing lacks a broadly adopted, robust framework for false discovery rate (FDR) control: because a prediction is generated rather than selected from a fixed list of candidates, the target–decoy strategy used for FDR control ^9^ and the entrapment strategy used to evaluate FDR control ^10^ in database search do not transfer directly. Consequently, the reliability of any individual detection is harder to establish. Careful validation therefore depends heavily on manual inspection, for which two kinds of evidence are especially informative. The first is the annotated MS/MS spectrum, which illustrates how completely the predicted fragment ion series (such as b- and y-ions) are supported by the observed peaks. Here Casanovo’s amino acid-level confidence scores can be particularly valuable, indicating the location of locally ambiguous positions along the peptide. The second type of evidence is the relationship between each predicted peptide and the reference proteome. Mapping *de novo* peptides to known proteins helps distinguish predictions supported by an existing protein sequence from candidate sample-specific peptides that are absent from the reference proteome, such as sequence variants. Both steps are essential to turning raw predictions into interpretable biological evidence, yet neither is provided by the command-line tool, and each ordinarily requires separate software and manual file handling.

Graphical front-ends have repeatedly broadened the user base of otherwise expert-only proteomics software. ^11–13^ For *de novo* sequencing in particular, the DeNovoGUI application brought together several classical sequencing algorithms under a single graphical interface. ^14^ However, this interface predates and does not support the deep learning models such as Casanovo that now define the state of the art. In addition, several tools have been developed specifically to make Casanovo easier to run and its results easier to interpret ^15^: a Docker image and a Nextflow workflow run the sequencing step within a containerized pipeline, the PDV viewer renders annotated spectra, ^16^ and the Limelight web application supports browsing, quality control, and sharing of results. ^17^ Each of these tools, however, addresses only one part of the analysis workflow and must be installed and connected separately. Running Casanovo itself still requires command-line or workflow tooling. To our knowledge, no self-contained, cross-platform desktop application has previously integrated installation, the Casanovo functions, and result interpretation behind a single point-and-click interface.

Here we present CasanovoGUI, a cross-platform desktop application that makes all of Casanovo’s main analysis functions available through a graphical interface while automating the installation and dependency management that its command-line use otherwise requires. CasanovoGUI further extends the workflow beyond sequencing to the interpretation steps—spectrum inspection and mismatch-aware mapping of predicted peptides to a proteome—that turn raw predictions into biological conclusions.

## Methods

CasanovoGUI is written in Java, with a graphical interface built using JavaFX, and is distributed as native installers for Windows, macOS, and Linux, plus a platform-independent executable JAR that runs on any system with a Java runtime. The native installers are produced with the jpackage tool and bundle a Java runtime, so the user does not need to install Java. The application handles installation, command orchestration, live output streaming, and interactive, linked visualization of the results, while the core sequencing is carried out by the official Casanovo program.

The feature that most directly removes the barrier to entry is automatic, self-contained installation of Casanovo. The first time the user starts an analysis, if the Casanovo Python package cannot be found, then CasanovoGUI offers to install it. The software automatically downloads uv (https://github.com/astral-sh/uv), a standalone Python package manager, creates a private virtual environment with a pinned Python interpreter under a single directory in the user’s home folder, and installs Casanovo into it without requiring administrator privileges. Critically, the installer manages the scientific Python and GPU dependencies that frequently cause manual installations to fail: it detects the GPU and installs a matched PyTorch build (CUDA or CPU), pins the native-library versions whose incompatibilities most commonly crash command-line Casanovo, and selects a compatible compute device on each platform. The application can also drive an existing Casanovo executable or one inside a named Conda environment, allowing advanced users to run customized Casanovo versions and integrate the GUI with environments that they already maintain. CasanovoGUI was developed and tested with Casanovo v5.2.1 and is designed to remain compatible with subsequent releases. Casanovo runs on either a GPU or the CPU alone, so a GPU shortens a run but is not required. Run times and memory use measured on each device are reported in the repository documentation (https://github.com/Noble-Lab/CasanovoGUI) and updated for each major Casanovo release.

Beyond local execution, CasanovoGUI can offload the Casanovo computation to a remote Linux machine, so that a user whose local computer lacks a suitable GPU, or for whom CPU execution is too slow, can take advantage of a more capable machine, such as a lab GPU server or a rented cloud instance on Amazon EC2 or Microsoft Azure, without leaving the interface. After the remote machine is configured once, an analysis is launched exactly as it would be locally: CasanovoGUI transfers the input files, sets up and runs Casanovo on the remote machine, streams its live output and progress back, and retrieves the results when the run finishes.

CasanovoGUI additionally relies on three external programs, each downloaded automatically on first use so that users do not need to install them separately. Two support results interpretation and are launched with the bundled Java runtime: the PDV spectrum viewer (v2.6.0) ^16^ for annotated-spectrum visualization, and `pepmap` (https://github.com/wenbostar/pepmap, v2.0.0) for mismatch-tolerant peptide-to-protein mapping. The third, ThermoRawFileParser (v2.0.0-dev), ^18^ converts Thermo `.raw` input files to mzML before a run.

To illustrate CasanovoGUI’s interpretation features, we analyzed a publicly available proteomics dataset from the human Jurkat cell line. ^19^ This dataset was originally generated to detect variant peptides arising from nonsynonymous nucleotide differences, by searching the spectra against a customized protein database built from matched RNA-seq data of the same cells. We downloaded the MS/MS data from PeptideAtlas (accession PASS00215) and sequenced them with Casanovo (v5.2.1) using its default parameter settings.

## Results

### A point-and-click interface to Casanovo

CasanovoGUI presents each of Casanovo’s analysis functions as its own tab (Figure 1): *de novo* sequencing (`casanovo sequence`) from mzML, mzXML, or MGF spectrum files ^6^; database search (`casanovo db-search`) using Casanovo as the scoring function against a protein FASTA file ^8^; evaluation of *de novo* predictions against annotated spectra (casanovo sequence --evaluate); and model training and fine-tuning (`casanovo train`). Each of these four analysis tabs corresponds directly to a single Casanovo command-line invocation, and each is run on its own rather than as one stage of a sequential workflow. A separate *View* tab provides post-run results interpretation, described below. CasanovoGUI also accepts vendor raw data directly as input to *de novo* sequencing and database search, so the user does not need to convert it before-hand. Thermo `.raw` files are converted to mzML with ThermoRawFileParser ^18^ before invoking Casanovo; as with the other helper tools, the converter is downloaded automatically on first use. Bruker `.d` (timsTOF) data are passed directly to Casanovo, which reads them natively. Inputs and outputs are chosen through file choosers and form fields rather than by typing paths, and model weights are optional—when left blank, the application detects and displays Casanovo’s cached default weights so the user knows which model will run. Casanovo has dozens of parameters that are normally set by editing a configuration file in YAML format.

**Figure 1:**
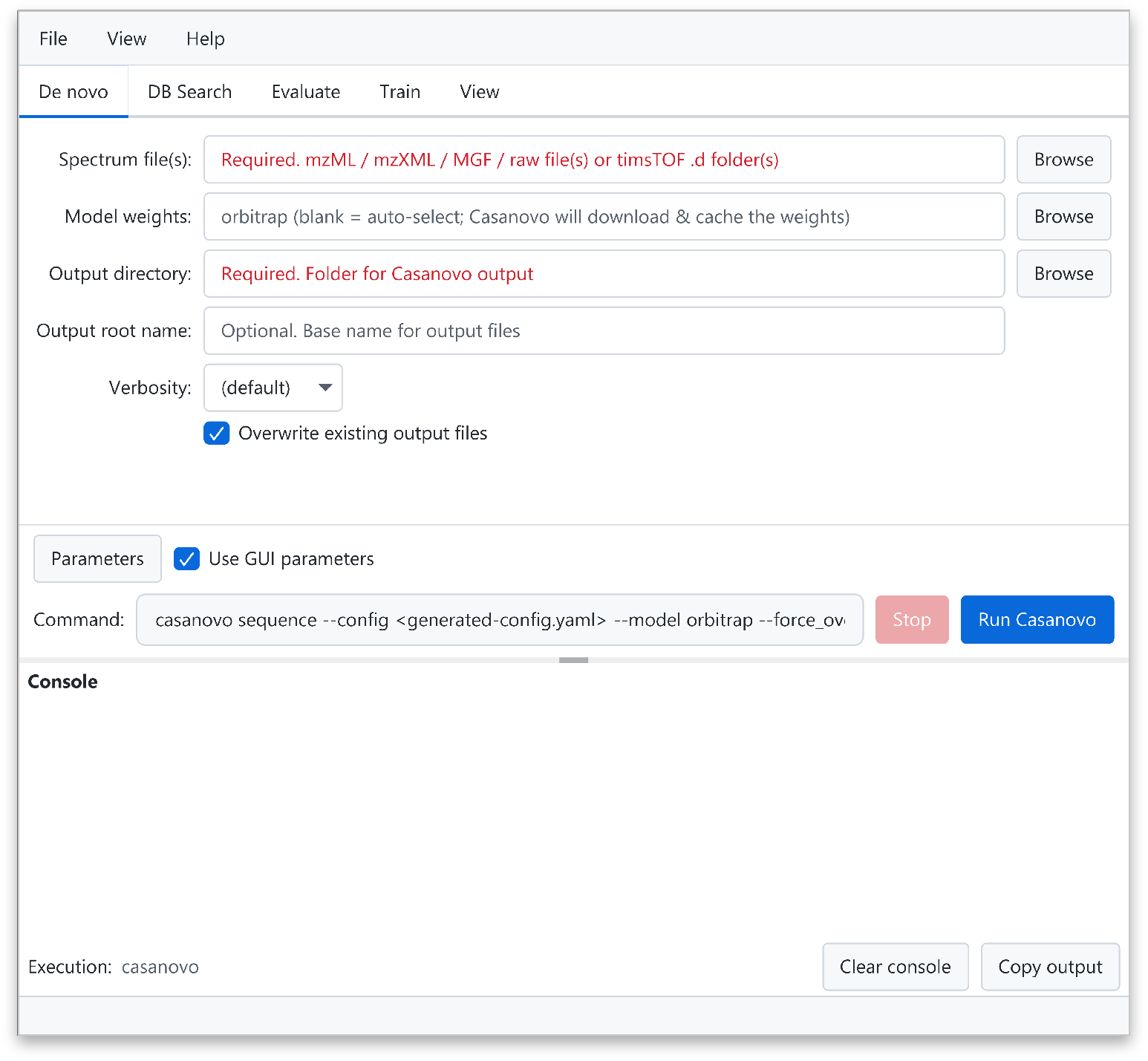
The CasanovoGUI interface, showing the *De novo* tab. A row of tabs selects the Casanovo function—*de novo* sequencing, database search, evaluation, or training—or the *View* tab for results interpretation. Inputs and outputs are chosen through form fields and file browsers; the *Parameters* button opens a dialog showing every configuration option, and the *Command* field previews the exact command line that will be executed. *Run Casanovo* starts the analysis and streams its output into the console below.

CasanovoGUI presents every parameter in a dialog organized into logical groups, each rendered with an editor appropriate to its type and annotated with a description drawn from Casanovo’s documentation, so a setting can be understood without consulting external references. The dialog can also export the assembled settings to a Casanovo YAML configuration file. Below the tabs, a read-only command preview always shows the exact command line that will be executed, keeping the interface transparent and reproducible from a terminal if desired. The exact configuration used for each run is saved alongside the results, providing a record of the parameters. Long-running analyses stream their output into a console panel in real time. CasanovoGUI interprets the updates emitted by the underlying tools and produces a progress bar derived from the spectrum count and batch size, giving feedback that the command line does not expose when its output is captured by another program.

### Mapping *de novo* peptides to a reference proteome

*De novo* sequencing produces predicted peptide sequences without directly linking them to protein origins. Mapping these peptides to a reference proteome provides important biological context by distinguishing predictions that match known protein sequences exactly, those that are compatible with known proteins only through one or more mismatches and may therefore represent sequence variants, and those that do not map to any known protein and are therefore absent from the reference proteome. Accordingly, CasanovoGUI includes a *View* tab that maps the peptides in a Casanovo result, reported in the mzTab format, ^20^ back onto a user-supplied protein FASTA database using `pepmap` (Figure 2). After a Casanovo run, sequencing results are loaded into the *View* tab automatically, so predicted peptides can be mapped by supplying a reference database and clicking *Run*, without re-selecting the results file. Internally, `pepmap` performs the mapping with the PeptideMapper algorithm^21^ from the compomics-utilities library, ^22^ which indexes the protein database with an FM-index ^23^ for fast lookup and supports mismatch-tolerant matching and protein regions containing the wildcard residue X—the one-letter code for an unspecified amino acid, which appears where a reference sequence leaves a position undetermined.

**Figure 2:**
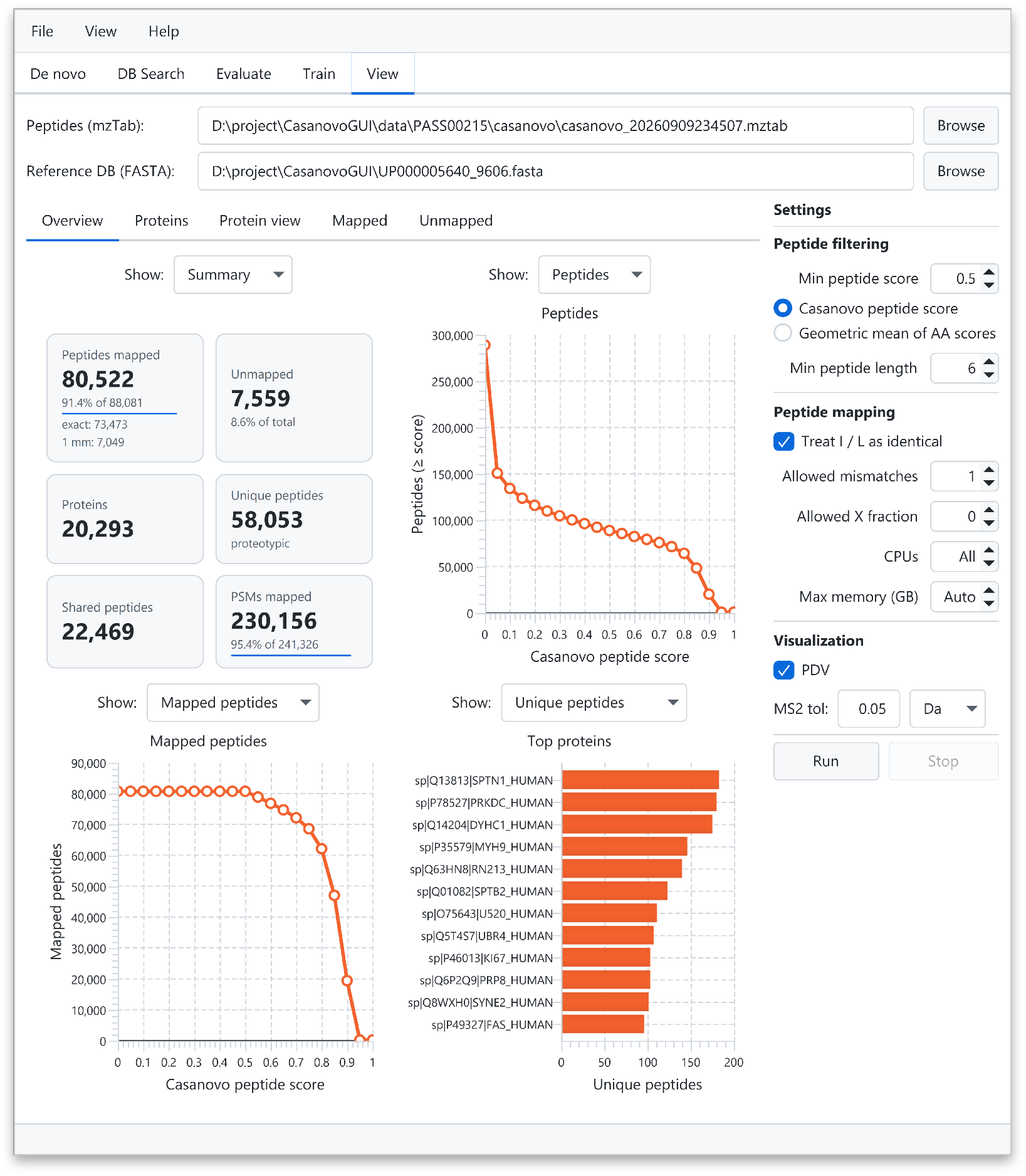
The integrated proteome-mapping view, showing the *Overview* tab of the *View* panel after mapping a Casanovo result against the human reference proteome with `pepmap`. The summary cards report the numbers of mapped peptides (with exact matches and mismatches counted separately), unmapped peptides, mapped proteins, unique and shared peptides, and mapped PSMs. The plots show the number of distinct peptides at or above each score (top right), the number of mapped peptides as a function of the score cutoff (bottom left), and the most peptide-rich proteins (bottom right). Mapping options are set in the panel at right.

Before mapping, the predicted peptides can be filtered by a minimum peptide length and a minimum peptide score. The score cutoff can be applied to either of two peptide-level scores: Casanovo’s own peptide score (`search_engine_score[1]` in the mzTab), which is the product of its per-residue confidence scores, or a length-normalized score, the geometric mean of the same per-residue scores. The mapping is governed by a few options: treating isoleucine and leucine as identical (the default, since they are indistinguishable by mass), the maximum number of allowed mismatches, and the maximum fraction of the aligned region that may consist of the wildcard residue X. Genomic variation is one of the major sources of sample-specific peptides that are absent from a reference proteome, in particular single amino acid variants. Thus, allowing a small number of mismatches during mapping is especially useful for interpreting them: a peptide that does not match the reference exactly but aligns to a known protein with a single substitution is a strong candidate for such a variant rather than a peptide that does not correspond to any reference protein. As a concrete example as shown in Figure 3, the *de novo* peptide SYLEVPLEENVNR maps to protein Q96CT7 with a single substitution, a tyrosine in place of the reference histidine. This same amino acid change was reported by earlier proteogenomic studies that searched a customized protein database derived from paired RNA-seq data, ^19,24^ independently corroborating the *de novo* sequencing detection.

**Figure 3:**
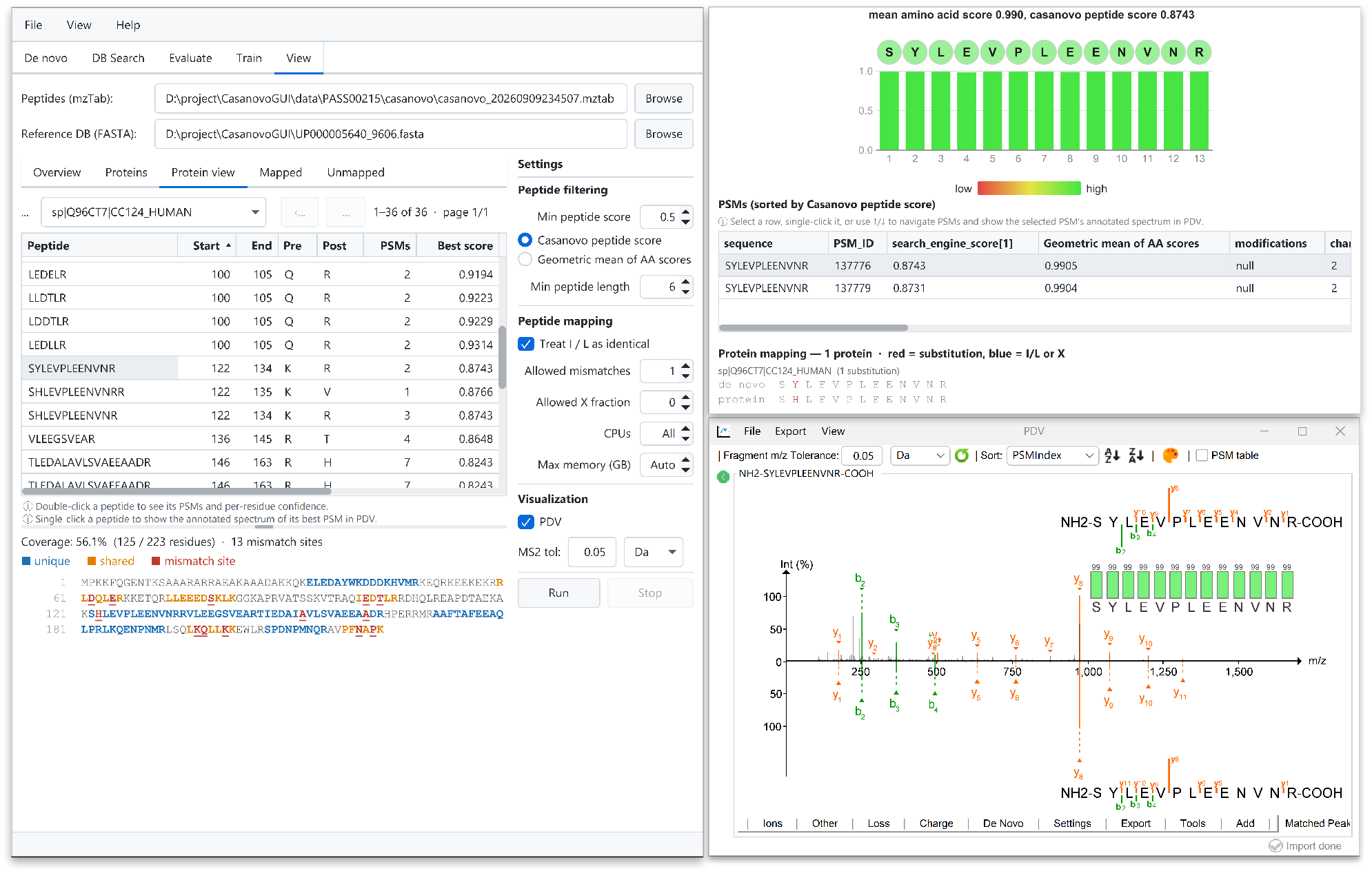
Linked, interactive visualization of *de novo* results across three windows driven from CasanovoGUI. *(Left)* The *Protein view* of the *View* tab lists the peptides mapped to a selected protein (here, Q96CT7) with their positions, PSM counts, and scores, and renders the protein’s sequence coverage by residue (blue, residues covered by unique peptides; orange, by shared peptides; red, mismatch sites where a mapped peptide differs from the protein). *(Top right)* Double-clicking a peptide opens the PSM table, which lists the peptide’s PSMs and displays the selected PSM’s amino acid-level scores as a colored bar track (green, high; red, low). Below the PSM table, a protein-mapping panel aligns the *de novo* peptide against each protein it maps to, marking substitution sites in red—here, the single tyrosine-for-histidine difference from Q96CT7—and isoleucine/leucine or wildcard-X positions in blue. *(Bottom right)* Selecting a PSM updates the linked PDV window to display the corresponding annotated observed spectrum—here for the peptide SYLEVPLEENVNR—with matched b and y ions and a per-residue confidence track. When predicted-spectrum display is enabled in PDV, a mirror plot compares the observed spectrum (above the axis) with the predicted spectrum (below the axis), generated here using AlphaPeptDeep through Koina.

The mapping results are presented as linked views. An *Overview* panel summarizes the mapped and unmapped peptides, the mapped proteins, and the numbers of unique peptides, which map to a single protein, and shared peptides, which map to more than one, together with plots of peptide counts versus score, the number of mapped peptides versus score cutoff, and the most peptide-rich proteins (Figure 2). *Proteins, Mapped*, and *Unmapped* tables list each protein with its peptide count, peptide–spectrum match (PSM) count, and sequence coverage, as well as each mapped peptide with its proteins, and each peptide that matched no protein—the last being precisely the peptides absent from the reference proteome that typically motivate *de novo* analysis. Selecting a protein in the *Proteins* table displays its coverage residue by residue, and clicking any peptide shows its best spectrum, so that a list of detections becomes a navigable, linked view of the underlying evidence. For proteins identified by a UniProt accession or entry name, hovering over the protein in the *Proteins* table retrieves its UniProt annotation on demand—protein name, gene, organism, length, and function—providing biological context without leaving the application.

### Visualizing and validating *de novo* results

Inspecting annotated spectra is an essential part of validating *de novo* results. After a sequencing run completes, CasanovoGUI stores the result and corresponding input spectra, so they can be opened in the PDV viewer from the *View* tab with no further file selection. A distinctive property of *de novo* sequencing is that confidence varies along the peptide: some residues are predicted with high confidence based on the observed fragment ions while others are not. Casanovo reports an amino acid level confidence score for every residue, which PDV renders as a color-coded track aligned with the peptide sequence in the annotated spectrum (Figure 3), shaded from green for high-confidence residues toward yellow and red as confidence decreases. This visualization allows a user to judge not merely whether a spectrum was sequenced, but which portions of the sequence are trustworthy—the assessment that manual validation of *de novo* results requires.

We also extended PDV with a control interface so that it can be driven interactively from CasanovoGUI. With a result open in PDV, clicking a peptide in any of the CasanovoGUI result tables selects that peptide’s best-scoring PSM and automatically renders its annotated spectrum in PDV, and the PSM table steps through the individual PSMs of a peptide as the user selects them—in each case without manual searching. CasanovoGUI communicates with PDV over the loopback interface (localhost), which lets two programs on the same computer exchange messages without those messages leaving the computer: selecting a PSM in CasanovoGUI sends the PSM’s identity to PDV, which responds by displaying the corresponding annotated spectrum. This keeps the two windows synchronized while they remain separate applications.

PDV can additionally present the observed spectrum and a reference spectrum together as a mirror plot, either to compare two candidate peptide interpretations of the same spectrum or to compare the observed spectrum against the fragmentation pattern predicted for the Casanovo peptide by a machine learning model such as Prosit, ^25^ UniSpec, ^26^ MS^2^PIP, ^27^ or AlphaPeptDeep. ^28^ These predicted fragment ion intensities are obtained on demand through the Koina prediction server. ^29^ The degree of agreement between the observed and predicted spectra is an additional, orthogonal indicator of the reliability of a Casanovo prediction. This visualization mode was introduced previously ^15^; here we extended it so that predicted mirror spectra can be generated automatically for every PSM rather than one at a time, and we also added support for several prediction models recently made available on the Koina server, letting the user choose among them.

Finally, CasanovoGUI can upload *de novo* results directly to Limelight^17^ for browser-based visualization and sharing with collaborators.

## Discussion

CasanovoGUI removes the principal practical barriers to using Casanovo by bringing its entire workflow— installation, all major analysis functions, and result interpretation—into a single, self-contained, cross-platform desktop application. A new user can go from download to a completed, inspected analysis without installing Java or Python, configuring a GPU stack, or composing a command line, and predictions can be assessed in place through annotated spectra, per-residue confidence scores, and mismatch-tolerant mapping to a reference proteome. These post hoc analysis capabilities are particularly important because *de novo* results lack a robust statistical error model. We anticipate that CasanovoGUI will be most valuable to experimentalists adopting *de novo* sequencing for the first time, to laboratories standardizing analyses across heterogeneous machines, and in teaching settings, while remaining useful to experienced users through its command transparency, Conda integration, and the option to run Casanovo on a remote machine.

### Data and software availability

CasanovoGUI is open source under the GPL-3.0 license. Source code, native installers for Windows, macOS, and Linux, and documentation are available at https://github.com/Noble-Lab/CasanovoGUI. Casanovo is available at https://github.com/Noble-Lab/casanovo, the PDV spectrum viewer (v2.6.0) at https://github.com/wenbostar/PDV, and the `pepmap` peptide- to-protein mapper (v2.0.0) at https://github.com/wenbostar/pepmap. The MS/MS data files used to generate the screenshots in Figures 2 and 3 were downloaded from the PeptideAtlas repository under accession PASS00215. ^19^

## Acknowledgments

This work was supported by the National Science Foundation award 2245300 (W.S.N.), the National Science Foundation Graduate Research Fellowship Program (Grant No. DGE-2140004, B.W.), National Institutes of Health grant R24GM141156, and the Research Foundation – Flanders (FWO G087625N). We gratefully acknowledge Yuanye Chi for testing CasanovoGUI. During the preparation of this manuscript, the authors used Claude (Anthropic) and ChatGPT (OpenAI) to assist with language editing and improve the clarity of the text. The authors reviewed and edited all AI-assisted content and take full responsibility for the final manuscript.

## Author contributions

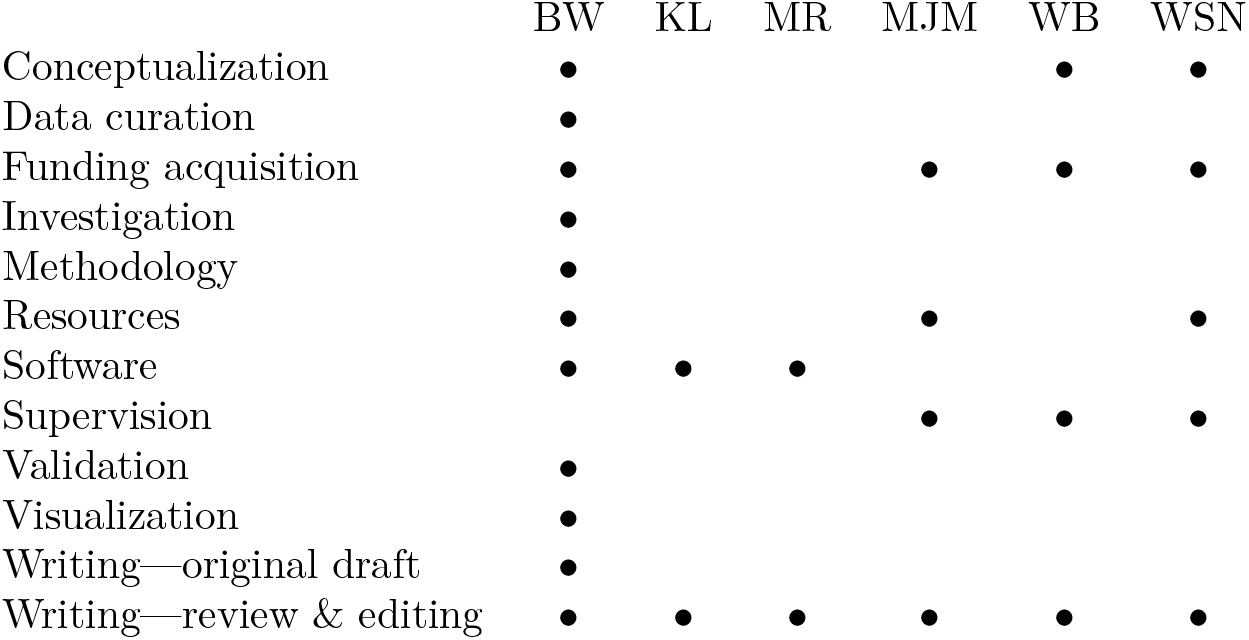

